# ggrain – a R package for raincloud plots

**DOI:** 10.1101/2024.12.13.628294

**Authors:** Nicholas Judd, Jordy van Langen, Davide Poggiali, Kirstie Whitaker, Tom Rhys Marshall, Micah Allen, Rogier Kievit

## Abstract

Clear data visualization is *essential* to effectively communicate empirical findings across various research fields. Raincloud plots fill this need by offering a transparent and statistically robust approach to data visualization. This is achieved by combining three plots in an aesthetically pleasing fashion. First, a dot plot displays raw data with minimal distortion, allowing a fast glance at the sample size and outlier identification. Next, a box plot displays key distributional summary statistics such as the median and interquartile range. Lastly, a violin plot transparently displays the underlying distribution of the data. Despite the widespread use of raincloud plots, an R-package in alignment with the *‘grammar of graphics’* was lacking. ‵ggrain‵ fills this need by offering one easy-to-use function (‵geom_rain‵) allowing the quick and seamless plotting of rainclouds in the R ecosystem. Further, it enables more complex plotting features such as factorial grouping, mapping with a secondary (continuous) covariate, and connecting observations longitudinally across multiple waves.

## Main Body

Data visualization is one of the most powerful and effective ways to communicate scientific findings. However, through force of habit or conventions, researchers commonly use visualization methods like the barplot, which obscures the distributional properties of underlying data. Bar plots are sensitive to distortion, unable to accurately represent the raw data, and do not display potential distribution differences^1^. For these reasons, they can lead to misinterpretation about the magnitudes of statistical differences between samples^2^ and are commonly criticized for being a non-transparent means to visualize data.

To overcome these challenges, we developed ‘raincloud plots’^1^, which address these problems in an intuitive, modular, and statistically robust format (Figure 1). This is achieved by combining three plots in an aesthetically pleasing fashion. First, a dot plot displays raw data with minimal distortion, allowing a fast glance at the sample size and outlier identification. Next, a box plot displays key distributional summary statistics such as the median and interquartile range. Lastly, a violin plot transparently displays the underlying distribution of the data. The combination of these plots allows maximal statistical information at a glance.

**Figure 1.**
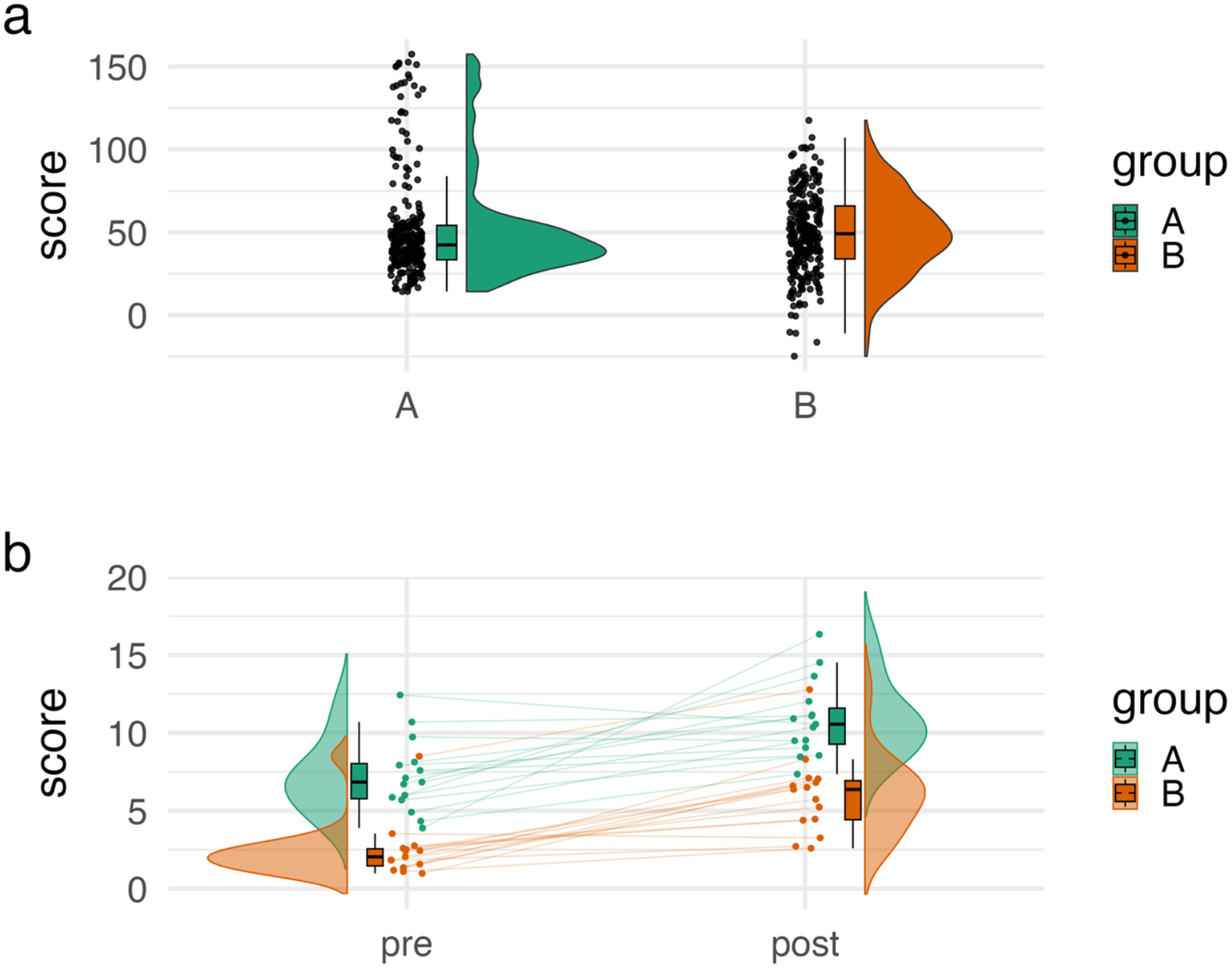
Two example figures using the ggrain package: **a)** illustrates distributional differences between two groups with raincloud plots, while **b)** shows intra-individual change following an intervention (pre/post) using a longitudinal raincloud subsetted by two groups. The source code is available at:https://github.com/njudd/ggrain/blob/main/inst/fig2.R.

Raincloud plots are well suited to illustrate statistical findings in pre/post or factorial grouped designs. For instance, to visualize the effect of an intervention in different mouse strains^3^, patient groups^4,5^, across cell types^6^, or policy changes^7^. One uniquely powerful aspect of raincloud plots is the ability to illustrate intraindividual change alongside average group change. This allows changes in variability (between subjects or groups) to be detected in longitudinal designs. For instance, in Figure 1b we can quickly see that the variability in scores between groups at the first timepoint is different (6.5 SD vs 2 SD), yet following an intervention the variability differences between groups disappear.

Despite the widespread use of raincloud plots, a package in alignment with the *‘grammar of graphics’* ^8–10^ in the R language was lacking. ‵ggrain‵ fills this need by offering one easy-to-use function (‵geom_rain‵) allowing the quick and seamless plotting of rainclouds in the R ecosystem; available on CRAN and Github (https://github.com/njudd/ggrain). Further, it enables more complex plotting features such as factorial grouping, mapping with a secondary (continuous) covariate, and connecting observations longitudinally across multiple occasions (Figure 1b). The full syntax to accomplish these more advanced features is outlined in our vignette (https://cran.r-project.org/package=ggrain/vignettes/ggrain.html).

Lastly, our group has implemented raincloud plots in other programming languages such as Python (https://pypi.org/project/ptitprince/) and JASP statistics (https://jasp-stats.org/)^11^. JASP offers an open-source GUI-based module to create raincloud plots – opening the door for researchers with limited programming experience to use raincloud plots and run statistical analyses.

Clear data visualization is *essential* to effectively communicate empirical findings across various research fields and the wider public. Raincloud plots and the ‵ggrain‵ package help facilitate intuitive, modular, and statistically robust data visualization.

## Acknowledgments

First, we would like to acknowledge the coining of the name ‘raincloud plots’ by Jon Roiser on March 15, 2018. We also would like to thank all participants in our in-person and online raincloud plots workshops, as their feedback has considerably improved our package.

## Ethics Approval

Not applicable, software package.

## Consent for publication

All authors give their consent for publication.

## Code availability

ggrain is an open-source R package available on Github (https://github.com/njudd/ggrain) and CRAN (https://cran.r-project.org/package=ggrain). As mentioned in the figure caption, the code and data for Figure 1 are on GitHub.

## References

1. Allen, M. et al. Wellcome open research 4, (2021).

2. Weissgerber, T.L., Milic, N.M., Winham, S.J. & Garovic, V.D. PLoS biology 13, e1002128 (2015).

3. Kimura, K. et al. Nat. Commun. 12, 3575 (2021).

4. Gil Avila, C. et al. eLife (2024). doi:10.7554/elife.101727.1

5. Tiddens, H.A.W.M. et al. Lancet Respir. Med. 10, 669–678 (2022).

6. Sasmita, A.O. et al. Nat. Neurosci. 27, 1668–1674 (2024).

7. Judd, N. & Kievit, R. bioRxiv 2025.01.17.633604 (2025). doi:10.1101/2025.01.17.6336048.

8. Wilkinson, L. (Springer: 2012).

9. Wickham, H. (Springer-Verlag New York: 2016).

10. Tiedemann, F. (2022). at <https://CRAN.R-project.org/package=gghalves>

11. Ott, V. L., van den Bergh, D., Boutin, B., van Doorn, J., Bartoš, F., Judd, N., … & Wagenmakers, E. J. (2024). Informative Data Visualization with Raincloud Plots in JASP (No. gv3ph_v1). Center for Open Science

